# Tracing the genetic origin of Europe’s first farmers reveals insights into their social organization

**DOI:** 10.1101/008664

**Authors:** Anna Szécsényi-Nagy, Guido Brandt, Victoria Keerl, János Jakucs, Wolfgang Haak, Sabine Möller-Rieker, Kitti Köhler, Balázs Gusztáv Mende, Marc Fecher, Krisztián Oross, Tibor Marton, Anett Osztás, Viktória Kiss, György Pálfi, Erika Molnár, Katalin Sebők, András Czene, Tibor Paluch, Mario Šlaus, Mario Novak, Nives Pećina-Šlaus, Brigitta Ősz, Vanda Voicsek, Krisztina Somogyi, Gábor Tóth, Bernd Kromer, Eszter Bánffy, Kurt W. Alt

## Abstract

Farming was established in Central Europe by the Linearbandkeramik culture (LBK), a well-investigated archaeological horizon, which emerged in the Carpathian Basin, in today’s Hungary. However, the genetic background of the LBK genesis has not been revealed yet. Here we present 9 Y chromosomal and 84 mitochondrial DNA profiles from Mesolithic, Neolithic Starčevo and LBK sites (7^th^/6^th^ millennium BC) from the Carpathian Basin and south-eastern Europe. We detect genetic continuity of both maternal and paternal elements during the initial spread of agriculture, and confirm the substantial genetic impact of early farming south-eastern European and Carpathian Basin cultures on Central European populations of the 6^th^-4^th^ millennium BC. Our comprehensive Y chromosomal and mitochondrial DNA population genetic analyses demonstrate a clear affinity of the early farmers to the modern Near East and Caucasus, tracing the expansion from that region through south-eastern Europe and the Carpathian Basin into Central Europe. Our results also reveal contrasting patterns for male and female genetic diversity in the European Neolithic, suggesting patrilineal descent system and patrilocal residential rules among the early farmers.

**Author Summary:** We report an exceptional large Neolithic DNA dataset from the Carpathian Basin, which was the cradle of the first Central European farming culture, the so called Linearbandkeramik culture. We generated 9 Y chromosomal and 84 mitochondrial DNA profiles from Mesolithic and Neolithic specimens from western Hungary and Croatia, attributed to the hunter-gatherers, Starčevo and LBK cultures (7^th^/6^th^ millennium BC). We observe genetic discontinuity between Mesolithic foragers and early farmers, and genetic continuity between farming populations of the 6^th^-4^th^ millennium BC across a vast territory of southeastern and Central Europe. Nine novel Y chromosome DNA profiles offer first insights into the Y chromosome diversity of the earliest European farmers, and further support the migration (demic diffusion) from the Near East into Central Europe along the Continental route of Neolithisation. The joint analyses of the two uniparental genetic systems let us conclude that men and women had a similar roles in the Early Neolithic migration process but their dispersal patterns were determined by sex-specific rules.

## Introduction

Agriculture was first established in the Near Eastern Fertile Crescent after 10,000 BC and expanded from the Levant and Anatolia to south-eastern Europe [1]. Archaeological research has described the subsequent spread of Neolithic farming into (and throughout) Central and south-western Europe along two major and largely contemporaneous routes. On the Continental route, the Carpathian Basin connected south-eastern Europe to the Central European loess plains, while the Mediterranean route bridged the eastern and western Mediterranean coasts, introducing farming to the Iberian Peninsula in the far West [2–6].

On the Continental route, the Early Neolithic Starčevo culture (STA) has played a major role in the Neolithisation of south-eastern Europe. The STA expanded from present-day Serbia to the western part of the Carpathian Basin, encompassing the regions of today’s northern Croatia and south-western Hungary (ca. 6,000-5,400 BC) [7,8] (Figure 1), and resulting in the formation of the Linearbandkeramik culture (LBK) [9]. The earliest LBK emerged in the mid-6^th^ millennium BC in Transdanubia (called “LBK in Transdanubia”, or LBKT) [9], marking the beginning of sedentary life in northern Hungary and via this area, Central Europe. The earliest LBKT coexisted with the STA in Transdanubia for about 100-150 years [10]. Archaeological research described an interaction zone between indigenous hunter-gatherer groups and farmers at the northernmost extent of the STA in Transdanubia, which might have led to the genesis of the LBKT [10,11]. After its formative phase in western Hungary, the LBK spread rapidly to Central Europe, reaching central Germany around 5,500 BC [2,12]. In the following 500 years, the LBK continually expanded, eventually covering a vast geographic area from the Paris Basin to Ukraine in its latest phase [2,13], and persisted in Transdanubia until ∼4,900 BC (Figure 1).

**Figure 1.**
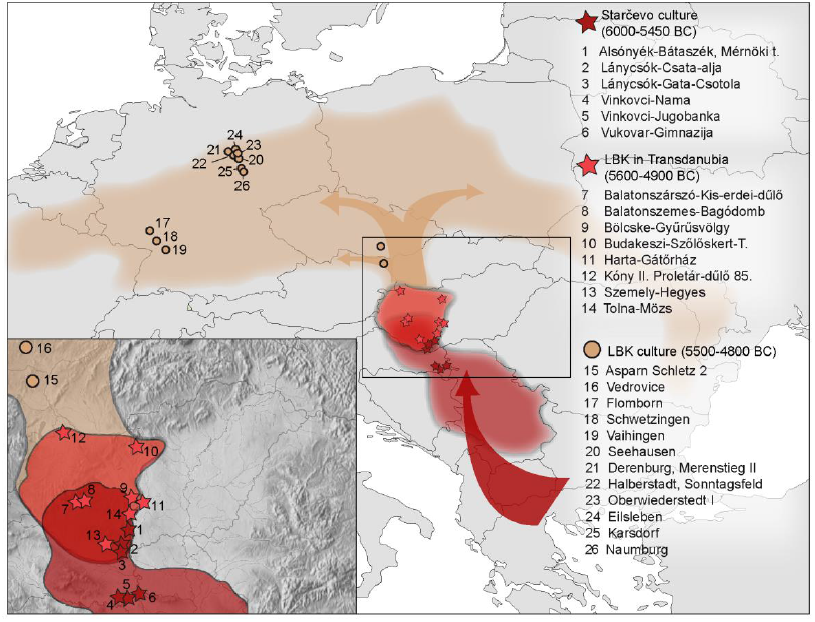
Geographic distribution of the Starčevo, LBK cultures and locations of the studied sites. The shaded areas of the maps show the distribution of the Starčevo culture (STA) and the LBK in Transdanubia (LBKT) and Central Europe (LBK) [2,3,5,7,9]. The arrows show the direction of the farmers’expansion into Central Europe, suggested by the above cited archaeological records. Coloured points indicate the studied sites of the STA (dark red) and LBKT (red) in western Hungary and northern Croatia. Central European LBK sites (brown) that were included in the comparative analyses are presented as well.

Despite the well-established archaeological relations between the STA and LBKT in the Carpathian Basin and the LBK in Central Europe, their genetic relationship has hitherto been unknown. Traditionally, scholars have explained the Neolithic transition either as an expansion of early farmers from the Near East, who brought new ideas as well as new genes (demic diffusion) [14–17], or as an adoption of farming technologies by indigenous hunter-gatherer populations with little or no genetic influence (cultural diffusion) [18–21]. These two contrasting models have been merged into complex integrationist approaches, considering small-scale population movements on regional levels [1,2,10,22].

Inferences drawn from genetic studies based on present-day data have yielded contradictory results about the Neolithic impact on the genetic diversity of modern Europeans, showing a disparity between mitochondrial DNA (mtDNA) and Y chromosomal patterns. Several Y chromosome studies supported the Neolithic demic diffusion model [17,23,24], while most mtDNA and some Y chromosomal studies have proposed a continuity of Upper Palaeolithic lineages [20,21,25,26]. The contrasting mtDNA and Y chromosomal evidence has been explained by differences in evolutionary scenarios, such as sex-biased migration [27].

Recent ancient DNA (aDNA) studies have provided direct insights into the mtDNA and autosomal diversity of hunter-gatherers in Europe [28–33] and the Central European LBK [33–37], describing a clear genetic discontinuity between local foragers and early farmers [28,31,36]. Comparative analyses with present-day populations have revealed Near Eastern affinities of the mitochondrial LBK ancestry supporting the demic diffusion model and population replacement at the beginning of the Neolithic period [36,37]. Data on Y chromosomal diversity in Neolithic Europe is still scarce. Beside the recently described first Mesolithic and Neolithic hunter-gatherer Y chromosomal data [33,38], Y chromosome data have been reported from a few LBK samples [36], the Tyrolean Iceman [39], the southwest European Neolithic [40,41], and from the Late Neolithic Central Germany [42,43].

The postulated Near Eastern origin of Central Europe’s LBK farmers has so far only been inferred from modern-day population data. The first ancient mtDNA data from early Near Eastern farmers has been reported recently [44], however the genetic diversity in the vast territory from the Fertile Crescent to Central Europe has been largely unexplored. Consequently, our aims were to i) study the genetic diversity of the early farming Carpathian Basin cultures from both the mtDNA and Y chromosome perspectives, ii) examine whether men and women had different demographic histories, iii) investigate the contribution of the STA to the genetic variability of the LBKT and LBK, iv) reveal the potential genetic origins of the first farmers in Eurasia, and v) to assess the role of the Continental route in the European Neolithic dispersal.

In this study we present 84 mtDNA and 9 Y chromosomal DNA data from Mesolithic (6,200-6,000 BC), and Neolithic specimens of the STA and LBKT from western Hungary and Croatia, spanning ∼900 years (ca. 5,800-4,900 BC) of Neolithic period. The population genetic analysis allowed detailed insight into the role of archaeological cultures from the Carpathian Basin in the spread of farming from the Near East.

## Results

### Mitochondrial DNA

Using well established aDNA methods (Material and Methods), we genotyped mtDNA variability by sequencing the hyper-variable segment I and II (HVS-I/II) and 22 single nucleotide polymorphisms (SNPs) on the coding region of the mitochondrial genome [36]. Overall, we investigated 109 skeletons from one Mesolithic, six STA and eight LBKT sites from western Hungary and Croatia (Figure 1, Dataset S1-S2). We successfully genotyped endogenous HVS-I sequences of 84 individuals (hunter-gatherer=1, STA=44, and LBKT=39) yielding a success rate of 76% (Dataset S3). We also sequenced parts of HVS-II from 25 individuals with consistent and identical HVS-I motifs in order to increase the phylogenetic resolution and to detect potential intra-site maternal kinship. The analysis of haplogroup defining coding region SNPs provided reproducible profiles for 96 individuals, with a success rate of 86% (Dataset S3-S4).

The haplotype of the Mesolithic skeleton from the Croatian Island Korčula belongs to the mtDNA haplogroup U5b2a5 (Dataset S3). The sub-haplogroup U5b has been shown to be frequent in pre-Neolithic hunter-gatherer communities across Europe [28–30,32,33,45,46]. Contrary to the low mtDNA diversity reported from hunter-gatherers of Central/North Europe [28–30], we identify substantially higher variability in early farming communities of the Carpathian Basin including the haplogroups N1a, T1, T2, J, K, H, HV, V, W, X, U2, U3, U4, and U5a (Table 1). Previous studies have shown that haplogroups N1a, T2, J, K, HV, V, W and X are most characteristic for the Central European LBK and have described these haplogroups as the mitochondrial ‘Neolithic package’ that had reached Central Europe in the 6^th^ millennium BC [36,37]. Interestingly, most of these haplogroups show comparable frequencies between the STA, LBKT and LBK, comprising the majority of mtDNA variation in each culture (STA=86.36%, LBKT=61.54%, LBK=79.63%). In contrast, hunter-gatherer haplogroups are rare in the STA and both LBK groups (Table 1). Besides similar haplogroup compositions we also found comparable haplotype diversity values for each culture (STA=0.97674, LBKT=0.95277, LBK=0.95483).

**Table 1.**
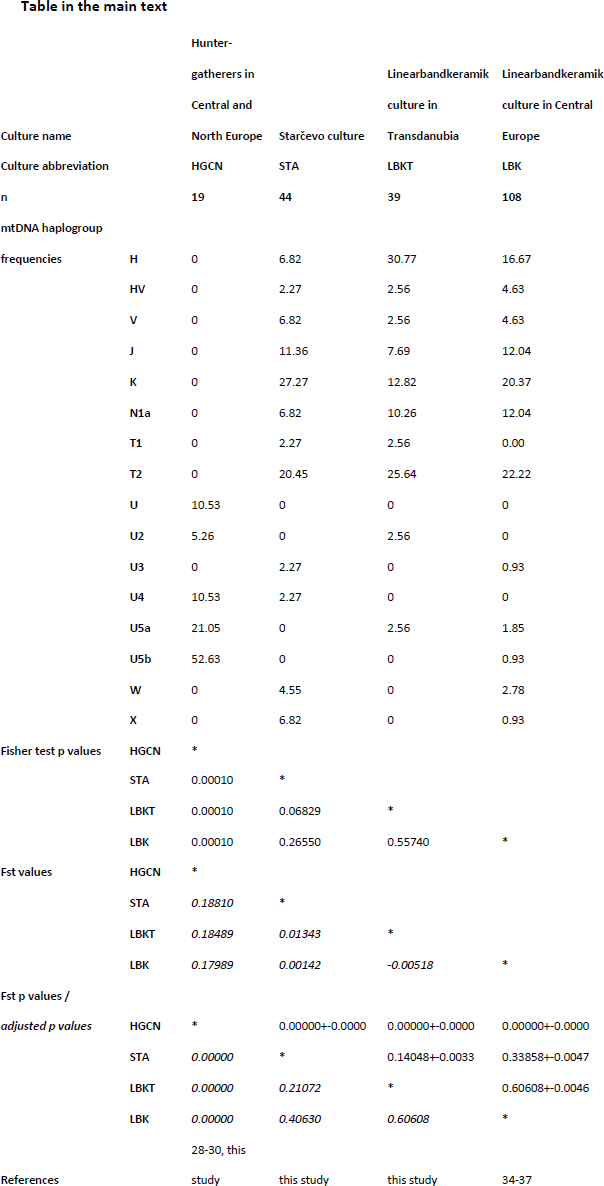
Mt haplogroup frequencies, Fisher’s exact test and genetic distances of four prehistoric metapopulations and archaeological cultures. Fisher’s exact test was based on mtDNA haplogroup frequencies. Genetic distances or Fst values (italicized) were calculated from HVS-I sequences (np 16056-16400). Genetic distance p values were post hoc adjusted to correct for multiple comparison by Benjamin and Hochberg method (italicized). Culture and population information are presented in Dataset S6.

In order to evaluate whether the haplogroup and haplotype composition of the STA, LBKT, LBK [34–37] and hunter-gatherers from Central/North Europe [28–30] differ significantly from each other, we performed a haplogroup-based Fisher’s exact test and a sequence based genetic distance analysis. In addition, we used the test of population continuity (TPC) [37], to elucidate whether the observed differences can be best explained by genetic drift or by other factors such as migration. These analyses reveal that the mtDNA composition of the Early Neolithic cultures is significantly different from that of the hunter-gatherers, both on the haplogroup (p=0.0001) and haplotype level (F_st_= 0.17989-0.18810, p=0.0000) (Table 1), indicating genetic discontinuity of maternal elements at the advent of farming in the Carpathian Basin as it has been reported previously from Central Europe [28,36,37]. The TPC shows that independent of the tested effective population size, the transition from hunter-gathering to farming cannot be explained by genetic drift alone (p<0.000001, Dataset S11). More importantly, non-significant differences between the haplogroup (p=0.06829-0.5574) and haplotype composition (F_st_=-0.00518-0.01343, p=0.21072-0.60608) of the STA and the LBK groups from Transdanubia and Central Europe (Table 1) support a rather homogenous mtDNA signature of early farming communities from both regions. The TPC also supports the scenario of population continuity during the Neolithic period, showing no significant p values among the pairwise compared Neolithic cultures (p>0.177 with all tested effective population sizes, Dataset S11).

We combined our Neolithic samples from the Carpathian Basin with 487 published mtDNA data from Upper Palaeolithic and Mesolithic [28–30,32,45,46], Neolithic [34–37,40,42,43,45–47] and Early Bronze Age [37] sites across Europe (Dataset S6) and conducted principal component analysis (PCA), multidimensional scaling (MDS), analysis of molecular variance (AMOVA) and shared haplotype analysis to compare the mtDNA variability of the STA and LBKT in a broader geographical and chronological context (Material and Methods).

PCA and MDS show that the mtDNA makeup of the STA and LBKT is strikingly similar to the LBK [34–37] and to subsequent cultures of the 5^th^/4^th^ millennium BC in Central Europe [37] (Figure 2-S1, Dataset S7-S8). This is predominately based on a high number of ‘Neolithic package’ lineages and low frequencies of haplogroups attributed to hunter-gatherers, which clearly distinguish this cluster from hunter-gatherers of Central/North [28–30] and southwest Europe [32,45,46], but also from Neolithic Iberian populations and Central European cultures of the 3^rd^/2^nd^ millennium BC (Figure 2). In order to exclude biases induced by potential maternal kinship within the prehistoric datasets, we performed PCA and MDS with a reduced dataset (*) as well, in which redundant haplotypes with identical HVS-I and II sequences from the same site were omitted. The reduced datasets have similar locations on the plots to the complete datasets, indicating that the effect of maternal kinship is negligible.

**Figure 2.**
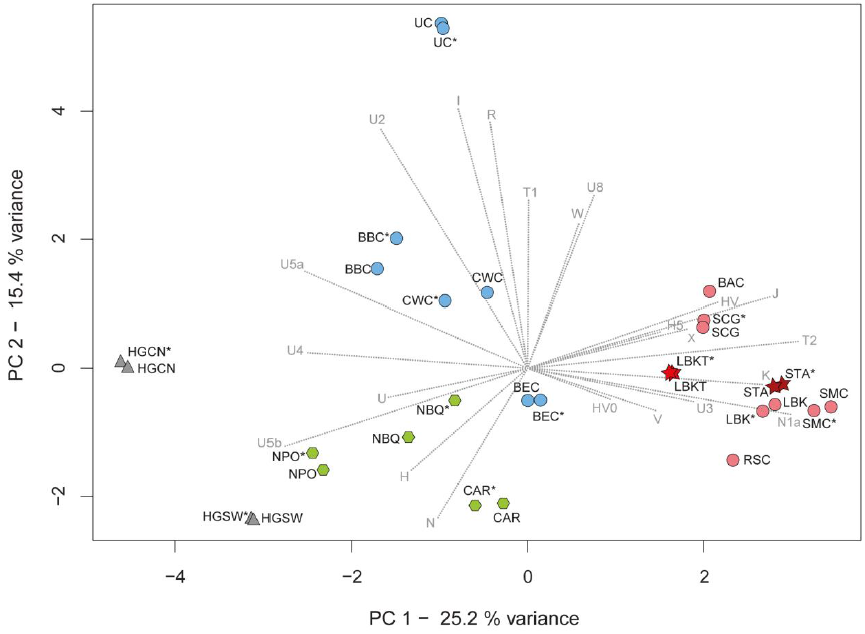
PCA plot comparing mtDNA data of 16 prehistoric cultures. PCA is based on the frequencies of 22 mtDNA haplogroups in the STA, LBKT and 14 prehistoric cultures. Color shadings and symbols denote cultures of different periods or European regions: hunter-gatherers (grey triangles), 6^th^ millennium BC Carpathian Basin cultures (red stars), LBK and 5^th^/4^th^ millennium BC cultures in Central Europe (rose circles), Central European 3^rd^/2^nd^ millennium BC cultures (blue circles), 6^th^/5^th^ millennium BC cultures of the Iberian Peninsula (green hexagons). The reduced version of each dataset is marked by an asterisk (*). The contribution of each haplogroup is superimposed as grey component loading vector. The first component (24.1%) clearly separates the hunter-gather populations and the Iberian cultures from the cluster of STA, LBKT, LBK and subsequent Central European 5^th^/4^th^ millennium BC cultures, while Central European cultures from the 3^rd^/2^nd^ millennium BC are differentiated along the second component (16.6%). Consequently, the PCA shows the maternal affiliation of the Carpathian Basin cultures to the LBK and to the 5^th^/4^th^ millennium BC cultures in Central Europe. Detailed information about the comparative data and haplogroups frequencies are listed in table S7. Culture abbreviations: hunter-gatherers in Central and North Europe (HGCN), hunter-gatherers in southwestern Europe (HGSW), Starčevo culture (STA), LBK in Transdanubia (LBKT), LBK in Central Europe (LBK), Rössen culture (RSC), Schöningen group (SCG), Baalberge culture (BAC), Salzmünde culture (SMC), Bernburg culture (BEC), Corded Ware culture (CWC), Bell Beaker culture (BBC), Únětice culture (UC), Cardial and Epicardial culture (CAR), Neolithic Basque Country and Navarre (NBQ), Neolithic Portugal (NPO).

We used AMOVA to evaluate whether the observed affinities of STA and LBKT with the LBK and 5^th^/4^th^ millennium BC cultures from Central Europe are the result of a shared population structure. We pooled HVS-I sequences from the STA and LBKT and nine archaeological cultures from Central Europe ranging from the LBK to the Early Bronze Age [34–37,42,43] into different groups, and tested 82 different arrangements to identify the constellation with the highest among-group variance and simultaneously with low variation within the groups (Dataset S9). The highest among-group variance was observed when STA and LBKT were arranged in one group with the Central European LBK and with all 5^th^/4^th^ millennium BC cultures, while the 3^rd^/2^nd^ millennium BC cultures were separated in a second group (among-group variation=3.50%, F_st_=0.03501, p=0.00396; within-group variation=0.20%, F_st_=0.00203, p=0.31139, Dataset S9). These results suggest a common genetic structure of the 6^th^-4^th^ millennium BC cultures.

We used shared haplotype analysis [48] and modified this approach by accounting for the temporal succession of cultures (ancestral shared haplotype analysis -ASHA). This enabled us to ascribe mtDNA lineages to particular cultures or time periods according to their first appearance in the dataset in chronological order (Figure 3, Dataset S10), and to estimate the amount of ancestral lineages in each culture, potentially derived from hunter-gatherers, STA, LBKT, LBK or other subsequent cultures. The ASHA shows that ancestral hunter-gatherer lineages were rare in the STA (2.27%), LBKT (0%) and LBK (1.85%) as well as in 5^th^/4^th^ millennium BC cultures (0%) and became more common in Central Europe during the 3^rd^/2^nd^ millennium BC (2.86-11.76%) [37]. In contrast, we identified a high degree of ancestral STA lineages in all subsequent cultures (LBKT=61.54%, LBK=55.56%, 5^th^/4^th^ millennium BC=36.84-63.64%, 3^rd^/2^nd^ millennium BC=36.17-43.18%). The subsequent LBKT reveals a smaller distinctive influence on its successors, since only 12.96% of the LBK, 0-10.53% of the 5^th^/4^th^ millennium BC, and 0-3.19% of the 3^rd^/2^nd^ millennium BC cultures can be traced back to ancestral lineages first observed in the LBKT. The number of new ‘ancestral’ lineages is even lower in the LBK of Central Europe, with no effect on the 3^rd^/2^nd^ millennium BC cultures.

**Figure 3.**
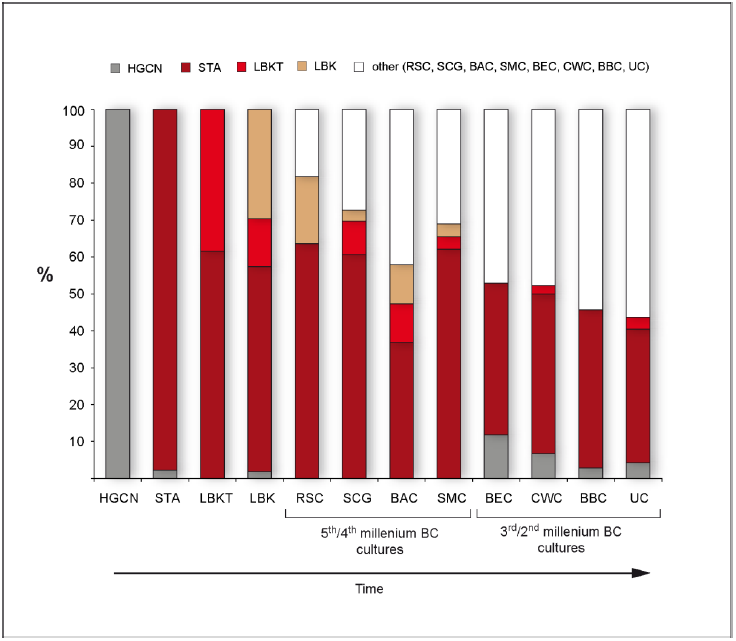
Results of the ancestral shared haplotype analysis. The bar plot shows the proportions of ancestral mtDNA lineages associated to hunter-gatherer (grey), STA (dark red), LBKT (red), LBK (brown), and other subsequent cultures (white) in the Central/North European hunter-gatherers dataset, the two Carpathian Basin cultures and nine Central European cultures ranging from the LBK to the Early Bronze Age. The mtDNA variability of the early Neolithic STA from western Carpathian Basin had a major influence on the mitochondrial gene pool of the Central European Neolithic cultures, which lasted from the LBK to the Early Bronze Age at least. Details of the ASHA are provided in table S10. Culture abbreviations: Hunter-gatherers in Central and North Europe (HGCN), Starčevo culture (STA), LBK in Transdanubia (LBKT), LBK in Central Europe (LBK), Rössen culture (RSC), Schöningen group (SCG), Baalberge culture (BAC), Salzmünde culture (SMC), Bernburg culture (BEC), Corded Ware culture (CWC), Bell Beaker culture (BBC), Únětice culture (UC).

In order to identify affinities of our Neolithic datasets with present-day populations, we collated 67,996 published HVS-I sequences from Eurasian populations and conducted PCA and genetic distance mapping (Material and Methods).

The PCA shows that the frequencies of N1a, T1, T2, K, J and HV, and the absence of Asian and African lineages in the Carpathian Basin cultures cause a clustering of the STA with populations of the Near East and the Caucasus, while the LBKT falls between the latter and populations from South and southeast Europe (Greeks, Bulgarians and Italians), which is caused by a higher frequency of haplogroup H in the LBKT (Figure S2, Dataset S12). However, the dominant frequencies of haplogroup N1a, T2, and K in the STA and LBKT result in a differentiation from all present-day populations along the third component.

Sequence-based genetic distance maps are largely consistent with PCA and reveal the greatest similarities of the STA to populations of the Near East (Iraq, Syria) and the Caucasus (Azerbaijan, Georgia, Armenia), as well as some European populations, such as Italy, Austria, Romania, and Macedonia (Figure 3a, Dataset S13). The distance map of the LBKT displays affinities that are overall similar to the STA, which includes populations from Azerbaijan, Syria, and Iraq. We also observe similarities to present-day Europeans, such as the populations of Great Britain, Portugal, Romania, Crete, and Russia (Figure 3b, Dataset S13). These similarity peaks are likely explained by elevated frequencies of shared lineages due to shared genetic drift in modern-day populations.

### Y chromosomal DNA

We also analysed the non-recombining part of the Y chromosome (NRY) in the investigated samples, using multiplex [36] and singleplex approaches, targeting 33 haplogroup defining SNPs. We successfully generated unambiguous NRY SNPs profiles for nine male individuals (STA=7, LBKT=2) (Dataset S3, S5). Three STA individuals belong to the NRY haplogroup F* (M89) and two specimens can be assigned to the G2a2b (S126) haplogroup, and one each to G2a (P15) and I2a1 (P37.2) (Dataset S3, S5). The two investigated LBKT samples carry haplogroups G2a2b (S126) and I1 (M253). Furthermore, the incomplete SNP profiles of eight specimens potentially belong to the same haplogroups; STA: three G2a2b (S126), two G2a (P15), and one I (M170); LBKT: one G2a2b (S126) and one F* (M89) (Dataset S5).

G2a2b and F* are rare in present-day Europe. Haplogroup G and its subgroups slightly increase towards the Near East and reach the highest frequency in populations of the south and northwest Caucasus [49,50], while haplogroup F* shows a diffuse dissemination pattern in Eurasia, which is based on insufficient sub-haplogroup resolution of most of the population genetic studies. Haplogroups I1 and I2a1 are most frequent in present-day populations of Europe, with the highest frequencies in Scandinavian [51–53] and southeast European populations respectively [51].

We used PCA and genetic distance maps to identify affinities of the Carpathian Basin samples with 49,516 NRY SNP profiles from present-day Eurasian and African populations (Material and Methods). Due to the similarities in Y chromosome composition and the small number of samples, we pooled STA and both LBK groups.

The elevated haplogroup G frequency in populations of the west Caucasus results in a clustering with the STA-LBK group on the second principal component the predominant frequencies of haplogroups G and F* lead to a clear separation of the STA-LBK group from all present-day populations along the third principal component (Figure S4, Dataset S14).

**Figure 4.**
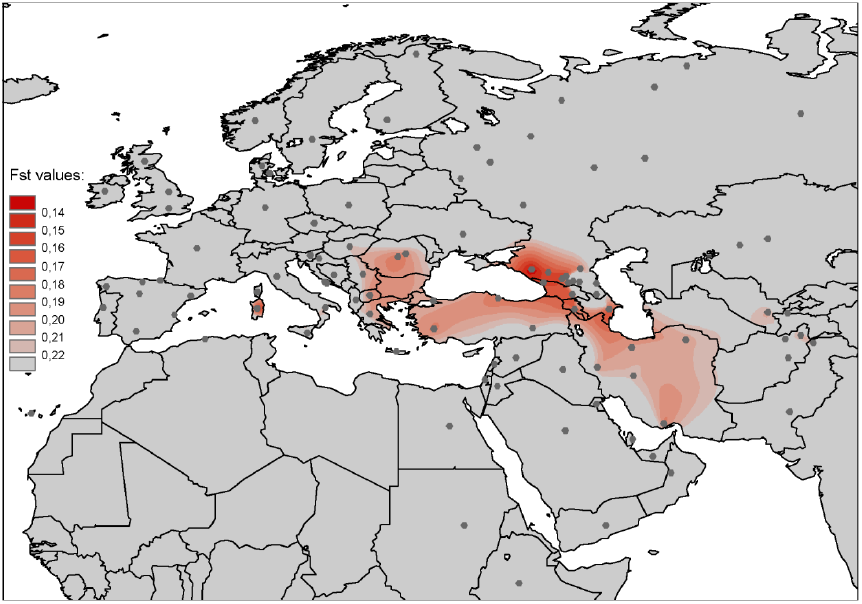
Genetic distance map of the STA-LBK Y chromosomal data. Y chromosomal genetic distances (F_st_) were computed between the STA-LBK samples and 100 present-day populations of Eurasia and North Africa and visualized on a geographic map. Grey dots denote the location of present-day populations. Color shadings indicate the degree of similarity or dissimilarity of Neolithic samples to the modern-day populations. Short distances and great similarities to present-day populations are marked by red areas. F_st_ values were scaled by an interval range of 0.01. F_st_ values higher than 0.21 were not differentiated (grey areas). The map shows remarkable affinities of the STA-LBK samples to present-day populations of the northwest and south Caucasus. Population information and F_st_ values are listed in table S15.

Similarly, the Y chromosome distance map discloses the greatest similarities to populations of the west and south Caucasus, such as Adyghe, Kabardin, Balkarians, Abkhazians, Azerbaijanis and Georgians as well as to the Sardinians (Figure 4, Dataset S15), which can be explained by the high frequency of haplogroup G/G2a [50,54] in these populations. This might reflect genetic drift, caused by isolation and small effective population size after a direct gene flow from the Near East, which lead to a fixation of this haplogroup [49]. Intriguingly, populations of the northeast Caucasus show greater distances to the STA-LBK samples due to lower abundance of haplogroup G/G2a [50]. Recently, the genomic data of an LBK individual from Stuttgart has been shown to be similar to modern-day Sardinians [33], which result can be explained by the isolation of the Sardinians, leading to the conservation of the Neolithic genetic signature. Nevertheless, our mtDNA population genetic analyses did not assure the Neolithic-Sardinian affinity, detected only on the NRY genetic distance map.

## Discussion

This study provides the first in-depth population survey of early farming cultures from the Carpathian Basin and south-eastern Europe and demonstrates their essential role in the genesis of the first farming communities of Central Europe. Our population genetic analyses (Fisher’s exact test, PCA, MDS, AMOVA, TPC) reveal a similar haplogroup composition and comparable haplotype diversity between the mtDNA variability of the Carpathian Basin cultures and the LBK from Central Europe (Table 1), indicating a homogenous and shared population structure of early farming communities from both regions (Figure 2, S1).

The ASHA shows that about 55% of the LBK lineages ascribed to characteristic ‘Neolithic package’ haplogroups could be traced back to the STA and LBK in Transdanubia (Figure 3, Dataset S10). It is therefore likely that this mtDNA signature was also present in ancient populations preceding the STA (7^th^/6^th^ millennium BC farming groups from the Aegean and the southern Balkans), in accordance with the archaeological record, which suggests cultural links to regions further southeast [5]. Interestingly, the STA mtDNA signature was still preserved in Neolithic cultures of the 5^th^/4^th^ millennium BC in Central Europe (Figure 3, Dataset S10), attesting a direct and enduring genetic legacy of the STA and LBKT in the Central European Neolithic, with minimal or no additional genetic influence from outside for the subsequent 2,500 years.

Importantly, our comparative analyses (PCA and genetic distance maps with modern population data) point out that both the mtDNA and NRY variability, observed in the Carpathian Basin samples, most likely originated in the Near East with connections to the Caucasus (Figure 4, S2-4), which is in accordance with previous mtDNA studies of the Central European LBK [36,37], and subsequent farming cultures of the 5^th^/4^th^ millennium BC [37]. The continuation of lineages through space and time suggests a scenario in which the genetic makeup of early farmers originated in the Near Eastern Fertile Crescent, from where it spread to Central Europe via the western Carpathian Basin, a region which acted as a natural corridor and an adaptation zone during the Neolithic expansion. The shared Near Eastern affinities of the STA, LBKT and LBK, and the genetic continuity in the maternal and paternal gene pools are consistent with the archaeological record, which describes the genesis of the early LBK (LBKT) from STA communities, followed by a rapid dispersal of the early LBK culture from Transdanubia towards the north-western part of Central Europe [3,9,13]. Recent aDNA study from 8000 BC Near Eastern farmers raises the question whether modern Near Eastern mtDNA can be used as a proxy for the Near Eastern Neolithic variability [44]. In our opinion, these newly described seven different incomplete HVS-I haplotypes (np 16095-16369) only provide a limited basis for comparative aDNA analyses, and we thus still consider modern-day Near Eastern genetic data sufficient proxies, when tracing the origin of the first European farmers. Recent study using ancient genomic data of the ‘Stuttgart’ LBK individual, the Tyrolean Iceman, and a Scandinavian farmer (Gök4) has shown rather south European than Near Eastern affinity of these ‘early farmers’, and has estimated a western hunter-gatherer ancestry of 0-45% in the early farmers’ gene pool [33]. These results do not contradict ours, since uniparental markers behave more conservative. They could preserve Near Eastern signature more consistently, even if admixture with foragers occurred on the way to Central Europe. Furthermore the results are not directly comparable with ours, since we had used earlier Neolithic specimens from a region that was nearer to the source region than it was the case in the study by Lazaridis et al.

The very low frequencies of hunter-gatherer lineages (0-2.27%), in the STA, LBKT and LBK sample sets (Figure 3) indicate that the arrival of agriculture in the Carpathian Basin and Central Europe was accompanied by a strong reduction of the currently known Mesolithic mtDNA substratum, resulting in a distinct and contrasting mtDNA haplogroup composition and significant differences between European hunter-gatherers and the Early Neolithic cultures (Figure 2-3, S1, Table 1, Dataset S7-8, S10-11). This scenario is consistent with coalescent-based simulations that have revealed genetic discontinuity between Central European hunter-gatherers and LBK communities [28,36]. The detection of haplogroup U5b in the investigated Mesolithic skeleton from Croatia matches previous observations, which describe sub-haplogroups of U as most frequent in forager populations across Europe, forming a characteristic Mesolithic mtDNA genetic substratum [28,37]. Residual Neolithic hunter-gatherer isolates, as reported from Central Europe by Bollongino et al. [30], have not yet been observed in our study region. According to the low proportion of hunter-gatherer mtDNA lineages in the LBK gene pool, we assume, that admixture between hunter-gatherers and colonizing LBK farmers was negligible in Central Europe. Considering the relative size and speed of the LBK expansion, we have to assume a substantial population growth during the earliest LBKT, which might have resulted in a population pressure and led to emigration from Transdanubia [55]. While such a radical population increase was not palpable from the Early Neolithic archaeological records [7], but recent extensive archaeological excavations have provided new insights into large-scale early LBKT settlements in western Hungary [9,56,57], which suggest larger source communities for a possible colonization than previously assumed.

Y chromosomal population genetic studies of modern-day Europeans have proposed that I1 and I2a1 NRY haplogroups were present in Europe since the Late Upper Palaeolithic. This was based on consistently high divergence time estimates [51,58], suggesting an expansion from Franco-Cantabrian (I1) and southeast European glacial refugia (I2a1) after the Last Glacial Maximum [51]. I2a1 has been recently described in Mesolithic specimens from Loschbour (Luxemburg) and Motala (Sweden) [33], in a Scandinavian Neolithic hunter-gatherer from Ajvide (Sweden, 2,900-2,600 BC) [38], as well as in Neolithic remains of southern France and northern Spain [40,41]. From the Mesolithic Motola site a further three men could be assigned to the haplogroup I [33]. The fact that almost all Mesolithic males belong to haplogroup I suggests that this haplogroup might represent a pre-farming legacy of the NRY variation in Europe.

Y chromosome haplogroups from STA and LBKT samples, such as haplogroups G2a2b and F* have also been reported from the Central European LBK [36], and support a close genetic relationship of the paternal lineages. Genetic studies on modern-day populations have discussed haplogroup G [25,59] and its subgroup G2a as potential representatives of the spread of farming from the Near East to Europe [26]. This scenario has recently been supported by Neolithic data from northern Spain [40] and southern France [41], which attested G2a a pivotal role in the Neolithic expansion on the Mediterranean route. Furthermore, G2a has also been reported from the Tyrolean Iceman (G2a2a1b (L91)) [39]. Taken together, these findings suggest that sub-haplogroups of G2a were frequent in Neolithic populations of the 6^th^-4^th^ millennia BC across Europe. Thus, if we take Y chromosomal haplogroup I2a (and possibly I1) as proxy for a Mesolithic paternal genetic substratum in Europe, we observe a similar pattern to the changeover in the mitochondrial DNA variability, in which NRY G lineages dominate Neolithic populations across Europe and I lineages become rare [36,39–43].

The most characteristic mtDNA haplogroup of early farmers from the Carpathian Basin and Central Europe is N1a. N1a has previously been discussed as a potential marker of the spread of farming [34]. The presence of N1a in early farmers from the Carpathian Basin (6.82-10.26%) and Central Europe (12.04%, Table 1) lends further support to its pivotal role as a marker for the Continental route of the Neolithic expansion. On the other hand, mtDNA N1a and NRY G2a haplogroups are rare in present-day European populations, which is also reflected in the separation of the 6^th^ millennium BC cultures from all present-day populations along the third principal component on the PCA plots (Figure S2, S4). These findings indicate further demographic events after the Early/Middle Neolithic period that shaped modern-day mtDNA and NRY variability. Recent evidence from ancient mtDNA has described the formation of modern-day variability by several successive migration events in Central Europe during the 3^rd^/2^nd^ millennium BC [37]. It is highly likely that these events have also affected the NRY diversity. Surprisingly, Y chromosome haplogroups, such as E1b1b1 (M35), E1b1b1a1 (M78), E1b1b1b2a (M123), J2 (M172), J1 (M267), and R1b1a2 (M269), which were claimed to be associated with the Neolithic expansion [23–25], have not been found so far in the 6^th^ millennium BC of the Carpathian Basin and Central Europe. Intriguingly, R1a and R1b, which represent the most frequent European Y chromosome haplogroups today, have been reported from cultures that emerged in Central Europe during the 3^rd^/2^nd^ millennium BC, while a basal R type has been reported from a Palaeolithic sample in Siberia [60] in agreement with a proposed Central Asian/Siberian origin of this lineage. In contrast, G2a has not been detected yet in late Neolithic cultures [42,43]. This suggests further demographic events in later Neolithic or post-Neolithic periods. However, we caution that the NRY record is still very small, especially in more recent periods, and further ancient Y data are required to shed light on the formation of the modern-day paternal diversity.

Interestingly, recent model-based statistical analyses of contemporary NRY and mtDNA data, testing a series of population scenarios for the Neolithic transition, have revealed a shared admixture history for men and women, but not the same demographic history [61]. This study has shown that female had a larger effective population size, likely based on differential effects of social and cultural practices including increasing sedentism alongside a shift to monogamy and patrilocality in early farmers. It is therefore important to interpret our new genetic data in the light of those findings. Considering the entire set of 32 published NRY records available for Neolithic Europe thus far, the low paternal diversity is indeed quite remarkable: G2a is the prevailing haplogroup in the Central European and Carpathian Basin Neolithic, and in French and Iberian Neolithic datasets [36,40,41]. There are only two exceptions, namely one E1b1b (V13) [41] individual from the Avellaner cave in Spain (∼5,000-4,500 BC), and two I2a [40] individuals from Treilles, France (∼3,000 BC). This very limited variation in NRY haplogroups in contrast to the high mtDNA haplogroups diversity suggests a larger effective population size for females than males. One plausible explanation for this phenomenon is patrilocality (where women move to their husband’s birth place after the marriage). Other possibilities that could lead to similar observations include polygyny or male-biased adult mortality. A patrilocal residential rule was possibly linked to a system of descent along the father’s line (patrilineality) in early farming communities. Ethnographic studies have suggested a change of residential rules at the advent of Neolithisation, showing different trends in residential rules among modern foragers and nonforagers [62]. Increasing sedentism promotes territorial defence and control of resources, favouring men in the inheritance of land and property, which consequently led to patrilocal residence [62]. At the same time, such residence pattern have to be momentarily flexible in expanding populations, allowing some of the sons to settle in new territories following population pressure and natural limitation of resources, e.g. after the carrying capacity of a particular region has been reached [61]. Patrilocality has also been raised in recent bioarchaeological studies. It has been suggested by aDNA evidence for the Treilles Neolithic community [41], and by stable isotope studies for the LBK in Central Europe [63].

It is important to note that patrilocality does not contradict the demic diffusion model, and it appears that both phenomena have left a discernible mark on the European Neolithic genetic diversity. While patrilocality and –lineality might have caused high mtDNA and low NRY within population diversity, the demic diffusion model best explains the mtDNA and NRY affinity of the early farmers to the modern Near East and Caucasus, and the observed global genetic homogeneity on a vast territory of south-eastern and Central Europe. Importantly, local processes of sex-biased migration are unlikely to have an effect on genetic variation at broader spatial scales. Our observations from many sites in Europe therefore argue for a common set of cultural and social practices across larger distances for early farming cultures in Europe. However, we caution that the observed differences in genetic diversity between males and females could also be influenced by resolution biases, resulting from the different sets of studied mtDNA and NRY markers. Examining sex-specific dynamics of early farmers is an important area that warrants further detailed research in order to address underlying parameters such as migration rate, level of exogamy and distances of marriage related dispersals among others.

The novel 83 mtDNA and nine NRY data from early farming Neolithic populations of the Carpathian Basin and one Mesolithic mtDNA profile help to fill the geographic gap on the Continental route of the Neolithic expansion from the Near Eastern Fertile Crescent to Central Europe. The joint analyses of mitochondrial and Y chromosomal DNA data support the demic diffusion of the early farmer men and women through western Hungary, and demonstrate the paramount importance of this region as a prehistoric corridor of the migration. We point out that archaeological cultures of the Carpathian Basin provided the genetic basis of the first Central European farmers that affected subsequent prehistoric cultures for a long period of time. Additionally, the new NRY data complement the sporadic European Y chromosomal dataset, and lend further support to patrilocal residential rules and patrilineal social system of the first farmers, underlining the role of demographic factors, which, depending strongly on cultural practices, notably shaped prehistoric and extant genetic diversity.

## Material and methods

We sampled one Mesolithic, 47 Starčevo and 61 LBKT skeletons, excavated in Croatia and western Hungary. The ancient DNA work was carried out in the Institute of Anthropology at the Johannes Gutenberg University of Mainz, following well-established protocols [34,36,37,42]. For minor modifications in the procedure of HVS-I, II and coding region SNP typing of the mitochondrial genome and SNP typing of the Y chromosome see Supplementary Information and Dataset S2-4.

The achieved genetic results were evaluated by population genetic analyses, using comparative ancient and modern DNA datasets (see Dataset S6, S12-15). We performed Fst and AMOVA analyses in Arlequin 3.5.1 [48]. Furthermore, we conducted Fisher’s exact test, PCA and MDS in R software environment. PCAs were based on mtDNA and Y chromosome haplogroup frequencies (Dataset S7, S12, S14). MDS was based on Slatkin linearized Fst values, calculated from mitochondrial HVS-I sequences (Dataset S8). Haplotype diversity was computed in DnaSP software, in version 5.10.01 [64].

MtDNA HVS-I sequence data and Y chromosomal haplogroup frequencies were applied for the genetic distance calculation, comparing the STA, LBKT mitochondrial DNA and a combined STA-LBK Y chromosomal datasets with 130 and 100 modern populations respectively (Dataset S13, S15). Genetic distance maps from the Fst values were generated in ArcGis version 10.0. In the ASHA that is a modified approach of the shared haplotype analysis [48], each HVS-I lineage within a given cultural dataset was traced back to its earliest appearance in a defined chronological order of the studied cultures. Each was regarded either as ancestral or as a new lineage, receiving its name after the culture where it was detected earliest in time (Dataset S10, Figure 3).

The adjusted parameters of the TPC [37] (Dataset S11) as well as further points of each analysis are detailed in the Supplementary Information.

## Acknowledgement

We thank Rozália Kustár, Olga Vajda-Kiss for archaeological information, Zsuzsanna K. Zoffmann for providing anthropological information and material, and Gabor Krizsma for informatics support.

## Author contributions

K.W.A., E.B., G.B., A.S-N. and J.J. designed the study; A.S-N. performed the palaeogenetic analyses; A.S-N., J.J., V.Ke., M.BG. and M.F. collected the samples; G.B. and A.S-N., collected reference data for the population genetic analyses; S.M-R. and A. S-N. cloned the PCR products; A.S-N., G.B., W.H. performed the biostatistics analyses; K.K., M. BG., B.Ő., G.T., E.M., G.P., M.Š., M.N. and N.P-Š. accomplished the anthropological analyses including individualization; K.O., T.M., V.Ki., A.O., K.S., A.C., V.V. K.So. and T.P. excavated the sites as well as provided the samples and the archaeological information and support; K.O., T.M., J.J. and A.S-N. summarized and reevaluated the archaeological data; B.K. made the radiocarbon dating; A.S-N., G.B., V.Ke., W.H. E.B. and K.W.A. wrote the paper; all authors discussed the results and commented on the manuscript.

## Financial Disclosure

This study is part of a 3-year project funded by the German Research Foundation (DFG) aimed at investigating the population dynamics of the Neolithic Carpathian Basin (AL 287-10-1). The funder had no role in study design, data collection and analysis, decision to publish, or preparation of the manuscript.

The project is a cooperation between the Bioarchaeometry Group of the Institute of Anthropology at the Johannes Gutenberg University of Mainz and the Institute of Archaeology, Research Centre for the Humanities, Hungarian Academy of Sciences in Budapest.

## Glossary

aDNA: Ancient DNA
AMOVA: Analysis of molecular variance ASHA Ancestral shared haplotype analysis
HVS I/II: Hyper Variable Segment I or II of the mitochondrial genome
LBK: Linearbandkeramik or Linear Pottery culture in Central Europe (refer to published LBK data from the Czech Republic, Lower Austria, and Germany)
LBKT: Linearbandkeramik or Linear Pottery culture in western Hungary/Transdanubia
MDS: Multidimensional scaling
mtDNA: Mitochondrial DNA
np: Nucleotide position
NRY: Non-recombining part of the Y chromosome PCA Principal component analysis
SNP: Single nucleotide polymorphism
STA: Starčevo culture
TPC: Test of population continuity

## References

1. Perlès C (2005) From the Near East to Greece: let’s reverse the focus – cultural elements that did not transfer. In: Lichter C, editor. BYZAS 2. How did farming reach Europe? Anatolian-European relations from the second half of the 7th through the first half of the 6th Millennium cal BC. Veröffentlichungen des Proceedings of the International Workshop Istanbul, 20-22 May 2004. Istanbul: Veröffentlichungen des Deutschen Archäologischen Instituts Istanbul. Ege Yayinlari. pp. 275–290.

2. Gronenborn D (1999) A variation on a basic theme : The transition to farming in southern Central Europe. J World Prehistory 13: 123–210.

3. Gronenborn D (2007) Beyond the models: “Neolithisation” in Central Europe. In: Whittle A, Cummings V, editors. Going over: the Mesolithic-Neolithic transition in North-West Europe. Oxford: Oxford University Press/British Academy. pp. 73–98.

4. Rowley-Conwy P (2011) Westward ho! The spread of agriculture from Central Europe to the Atlantic. Curr Anthropol 52: S431–S451. doi:10.1086/658368.

5. Price TD (editor) (2000) Europe’s first farmers. Price TD, editor Cambridge: Cambridge University Press. 429 p.

6. Zilhão J (2001) Radiocarbon evidence for maritime pioneer colonization at the origins of farming in west Mediterranean Europe. Proc Natl Acad Sci U S A 98: 14180–14185.

7. Kalicz N (2010) An Grenze “zweier Welten” -Transdanubien (Ungarn) im Frühneolithikum. In: Gronenborn D, Petrasch J, editors. Die Neolithisierung Mitteleuropas. The Spread of Neolithic to Central Europe. International Conference, 24-26th June 2005. Mainz RGZM. Mainz: RGZM. pp. 235–254.

8. Minichreiter K, Bronic IK (2006) New radiocarbon dates for the Early Starčevo Culture in Croatia. Pril Inst arh Zagreb 23: 5–16.

9. Oross K, Bánffy E (2009) Three successive waves of Neolithisation: LBK development in Transdanubia. Doc Praehist 36: 175–189. doi:10.4312/dp.36.11.

10. Bánffy E (2004) The 6th millennium BC boundary in western Transdanubia and its role in the Central European Neolithic transition (the Szentgyörgyvölgy-Pityerdomb settlement). Budapest: Archaeological Institute of the Hungarian Academy of Science. 451 p.

11. Bánffy E, Eichmann WJ, Marton T (2007) Mesolithic foragers and the spread of agriculture in western Hungary. In: Kozlowski JK, Nowak M, editors. Proceedings of the XV UISPP World Congress (Lisbon, 4–9 September 2006) Vol. 6. BAR IS 1726. Oxford. pp. 53–62.

12. Dolukhanov P, Shukurov A, Gronenborn D, Sokoloff D, Timofeev V, et al. (2005) The chronology of Neolithic dispersal in Central and Eastern Europe. J Archaeol Sci 32: 1441– 1458. doi:10.1016/j.jas.2005.03.021.

13. Bogucki P, Grygiel R (1993) The first farmers of Central Europe : a survey article. J F Archaeol 20: 399–426.

14. Ammerman AJ, Cavalli-Sforza LL (1984) The Neolithic transition and the genetics of populations in Europe. Princeton (New Jersey): Princeton University Press. 200 p.

15. Renfrew C (1987) Archaeology and Language. The Puzzle of Indo-European Origins. Cambridge: Cambridge University Press.

16. Pinhasi R, von Cramon-Taubadel N (2009) Craniometric data supports demic diffusion model for the spread of agriculture into Europe. PLoS One 4: e6747. doi:10.1371/journal.pone.0006747.

17. Chikhi L, Nichols RA, Barbujani G, Beaumont MA (2002) Y genetic data support the Neolithic demic diffusion model. Proc Natl Acad Sci U S A 99: 11008–11013. doi:10.1073/pnas.162158799.

18. Barker G (1985) Prehistoric farming in Europe. Cambridge: Cambridge University Press. 345 p.

19. Whittle A (1996) Europe in the Neolithic. Cambridge: Cambridge University Press. 459 p.

20. Richards M, Côrte-Real H, Forster P, Macaulay V, Wilkinson-Herbots H, et al. (1996) Paleolithic and neolithic lineages in the European mitochondrial gene pool. Am J Hum Genet 59: 185–203.

21. Richards M, Macaulay V, Hickey E, Vega E, Sykes B, et al. (2000) Tracing European founder lineages in the Near Eastern mtDNA pool. Am J Hum Genet 67: 1251–1276.

22. Zvelebil M (1989) On the transition to farming in Europe, or what was spreading with the Neolithic: a reply to Ammerman. Antiquity 63: 379–383.

23. Semino O, Magri C, Benuzzi G, Lin AA, Al-Zahery N, et al. (2004) Origin, diffusion, and differentiation of Y-chromosome haplogroups E and J: inferences on the neolithization of Europe and later migratory events in the Mediterranean area. Am J Hum Genet 74: 1023– 1034. doi:10.1086/386295.

24. Balaresque P, Bowden GR, Adams SM, Leung H-Y, King TE, et al. (2010) A predominantly neolithic origin for European paternal lineages. PLoS Biol 8: e1000285. doi:10.1371/journal.pbio.1000285.

25. Semino O, Passarino G, Oefner PJ, Lin AA, Arbuzova S, et al. (2000) The genetic legacy of paleolithic Homo sapiens sapiens in extant Europeans: a Y chromosome perspective. Science 290: 1155–1159. doi:10.1126/science.290.5494.1155.

26. Battaglia V, Fornarino S, Al-Zahery N, Olivieri A, Pala M, et al. (2009) Y-chromosomal evidence of the cultural diffusion of agriculture in Southeast Europe. Eur J Hum Genet 17: 820–830. doi:10.1038/ejhg.2008.249.

27. Rasteiro R, Bouttier P-A, Sousa VC, Chikhi L (2012) Investigating sex-biased migration during the Neolithic transition in Europe, using an explicit spatial simulation framework. Proc Biol Sci 279: 2409–2416. doi:10.1098/rspb.2011.2323.

28. Bramanti B, Thomas MG, Haak W, Unterlaender M, Jores P, et al. (2009) Genetic discontinuity between local hunter-gatherers and central Europe’s first farmers. Science 326: 137–140. doi:10.1126/science.1176869.

29. Fu Q, Mittnik A, Johnson PLF, Bos K, Lari M, et al. (2013) A revised timescale for human evolution based on ancient mitochondrial genomes. Curr Biol 23: 1–7. doi:10.1016/j.cub.2013.02.044.

30. Bollongino R, Nehlich O, Richards MP, Orschiedt J, Thomas MG, et al. (2013) 2000 years of parallel societies in Stone Age Central Europe. Science 342: 479–481. doi:10.1126/science.1245049.

31. Skoglund P, Malmström H, Raghavan M, Storå J, Hall P, et al. (2012) Origins and genetic legacy of Neolithic farmers and hunter-gatherers in Europe. Science 336: 466–469. doi:10.1126/science.1216304.

32. Sánchez-Quinto F, Schroeder H, Ramirez O, Avila-Arcos MC, Pybus M, et al. (2012) Genomic affinities of two 7,000-year-old Iberian hunter-gatherers. Curr Biol 22: 1494–1499. doi:10.1016/j.cub.2012.06.005.

33. Lazaridis I, Patterson N, Mittnik A, Renaud G, Mallick S, et al. (2014) Ancient human genomes suggest three ancestral populations for present-day Europeans. BioArxiv. http://dx.doi.org/10.1101/001552.

34. Haak W, Forster P, Bramanti B, Matsumura S, Brandt G, et al. (2005) Ancient DNA from the first European farmers in 7500-year-old Neolithic sites. Science 310: 1016–1018. doi:10.1126/science.1118725.

35. Bramanti B (2008) Ancient DNA: genetic analysis of aDNA from sixteen skeletons of the Vedrovice. Anthropol 46: 153–160.

36. Haak W, Balanovsky O, Sanchez JJ, Koshel S, Zaporozhchenko V, et al. (2010) Ancient DNA from European early Neolithic farmers reveals their Near Eastern affinities. PLoS Biol 8: e1000536. doi:10.1371/journal.pbio.1000536.

37. Brandt G, Haak W, Adler CJ, Roth C, Szécsényi-Nagy A, et al. (2013) Ancient DNA reveals key stages in the formation of Central European mitochondrial genetic diversity. Science 342: 257–261. doi:10.1126/science.1241844.

38. Skoglund P, Malmström H, Omrak A, Raghavan M, Valdiosera C, et al. (2014) Genomic diversity and admixture differs for Stone-age Scandinavian foragers and farmers. Science 344: 747–750. doi:10.1126/science.1253448.

39. Keller A, Graefen A, Ball M, Matzas M, Boisguerin V, et al. (2012) New insights into the Tyrolean Iceman’s origin and phenotype as inferred by whole-genome sequencing. Nat Commun 3: 698. doi:10.1038/ncomms1701.

40. Lacan M, Keyser C, Ricaut F-X, Brucato N, Tarrús J, et al. (2011) Ancient DNA suggests the leading role played by men in the Neolithic dissemination. Proc Natl Acad Sci U S A 108: 18255–18259. doi:10.1073/pnas.1113061108.

41. Lacan M, Keyser C, Ricaut F-X, Brucato N, Duranthon F, et al. (2011) Ancient DNA reveals male diffusion through the Neolithic Mediterranean route. Proc Natl Acad Sci U S A 108: 9788–9791. doi:10.1073/pnas.1100723108.

42. Haak W, Brandt G, de Jong HN, Meyer C, Ganslmeier R, et al. (2008) Ancient DNA, Strontium isotopes, and osteological analyses shed light on social and kinship organization of the Later Stone Age. Proc Natl Acad Sci U S A 105: 18226–18231. doi:10.1073/pnas.0807592105.

43. Lee EJ, Makarewicz C, Renneberg R, Harder M, Krause-Kyora B, et al. (2012) Emerging genetic patterns of the European Neolithic: perspectives from a late Neolithic Bell Beaker burial site in Germany. Am J Phys Anthropol 148: 571–579. doi:10.1002/ajpa.22074.

44. Fernández E, Pérez-Pérez A, Gamba C, Prats E, Cuesta P, et al. (2014) Ancient DNA Analysis of 8000 B.C. Near Eastern Farmers Supports an Early Neolithic Pioneer Maritime Colonization of Mainland Europe through Cyprus and the Aegean Islands. PLoS Genet 10: e1004401. doi:10.1371/journal.pgen.1004401.

45. Chandler H, Sykes B, Zilhão J (2005) Using ancient DNA to examine genetic continuity at the Mesolithic-Neolithic transition in Portugal. In: Arias P, Ontañón R, García-Moncó C, editors. Actas dell III Congreso del Neolítico en la Península Ibérica, Santander, Monografías del Instituto internacional de Investigaciones Prehistóricas de Cantabria 1. Vol. 1. pp. 781–786.

46. Hervella M, Izagirre N, Alonso S, Fregel R, Alonso A, et al. (2012) Ancient DNA from hunter-gatherer and farmer groups from Northern Spain supports a random dispersion model for the Neolithic expansion into Europe. PLoS One 7: e34417. doi:10.1371/journal.pone.0034417.

47. Gamba C, Fernández E, Tirado M, Deguilloux MF, Pemonge MH, et al. (2012) Ancient DNA from an Early Neolithic Iberian population supports a pioneer colonization by first farmers. Mol Ecol 21: 45–56. doi:10.1111/j.1365-294X.2011.05361.x.

48. Excoffier L, Lischer HEL (2010) Arlequin suite ver 3.5: a new series of programs to perform population genetics analyses under Linux and Windows. Mol Ecol Resour 10: 564–567. doi:10.1111/j.1755-0998.2010.02847.x.

49. Balanovsky O, Dibirova K, Dybo A, Mudrak O, Frolova S, et al. (2011) Parallel evolution of genes and languages in the Caucasus region. Mol Biol Evol 28: 2905–2920. doi:10.1093/molbev/msr126.

50. Yunusbayev B, Metspalu M, Järve M, Kutuev I, Rootsi S, et al. (2012) The Caucasus as an asymmetric semipermeable barrier to ancient human migrations. Mol Biol Evol 29: 359–365. doi:10.1093/molbev/msr221.

51. Rootsi S, Magri C, Kivisild T, Benuzzi G, Help H, et al. (2004) Phylogeography of Y-chromosome haplogroup I reveals distinct domains of prehistoric gene flow in Europe. Am J Hum Genet 75: 128–137. doi:10.1086/422196.

52. Karlsson AO, Wallerström T, Götherström A, Holmlund G (2006) Y-chromosome diversity in Sweden – A long-time perspective. Eur J Hum Genet 14: 963–970. doi:10.1038/sj.ejhg.5201651.

53. Lappalainen T, Laitinen V, Salmela E, Andersen P, Huoponen K, et al. (2008) Migration waves to the Baltic Sea region. Ann Hum Genet 72: 337–348. doi:10.1111/j.1469-1809.2007.00429.x.

54. Morelli L, Contu D, Santoni F, Whalen MB, Francalacci P, et al. (2010) A comparison of Y-chromosome variation in Sardinia and Anatolia is more consistent with cultural rather than demic diffusion of agriculture. PLoS One 5: e10419. doi:10.1371/journal.pone.0010419.

55. Petrasch J (2001) Seid fruchtbar und mehret euch und füllet die Erde und machet sie euch untertan. Archäologisches Korrespondenzblatt 31: 13–25.

56. Marton T, Oross K (2010) Siedlungsforschung in linienbandkeramischen Fundorten in Zentral- und Südtransdanubien-Wiege, Peripherie oder beides? In: Kreienbrink F, Cladders M, Stäuble H, Tischendorf T, Wolfram S, editors. Siedlungsstruktur und Kulturwandel in der Bandkeramik. Beiträge der internationalen Tagung “Neue Fragen zur Bandkeramik oder alles beim Alten?!” Leipzig, 23. bis 24. September 2010. Arbeits-und Forschungsberichte zur sächsischen Bodendenkmalpflege, Beihe. Dresden: Druckhaus Dresden GmbH. pp. 220–239.

57. Oross K, Marton T (2012) Neolithic burials of the Linearbandkeramik settlement at Balatonszárszó and their European context. Acta Archaeol 63: 257–299. doi:10.1556/AArch.63.2012.2.1.

58. Pericić M, Lauc LB, Klarić IM, Rootsi S, Janićijevic B, et al. (2005) High-resolution phylogenetic analysis of southeastern Europe traces major episodes of paternal gene flow among Slavic populations. Mol Biol Evol 22: 1964–1975. doi:10.1093/molbev/msi185.

59. Behar DM, Garrigan D, Kaplan ME, Mobasher Z, Rosengarten D, et al. (2004) Contrasting patterns of Y chromosome variation in Ashkenazi Jewish and host non-Jewish European populations. Hum Genet 114: 354–365. doi:10.1007/s00439-003-1073-7.

60. Raghavan M, Skoglund P, Graf KE, Metspalu M, Albrechtsen A, et al. (2014) Upper Palaeolithic Siberian genome reveals dual ancestry of Native Americans. Nature 505: 87–91. doi:10.1038/nature12736.

61. Rasteiro R, Chikhi L (2013) Female and male perspectives on the Neolithic transition in Europe: clues from ancient and modern genetic data. PLoS One 8: e60844. doi:10.1371/journal.pone.0060944.

62. Marlowe FW (2004) Marital residence among foragers. Curr Anthropol 45: 277–284. doi:10.1086/382256.

63. Bentley RA, Bickle P, Fibiger L, Nowell GM, Dale CW, et al. (2012) Community differentiation and kinship among Europe’s first farmers. Proc Natl Acad Sci U S A 109: 9326–9330. doi:10.1073/pnas.1113710109.

64. Librado P, Rozas J (2009) DnaSP v5: a software for comprehensive analysis of DNA polymorphism data. Bioinformatics 25: 1451–1452. doi:10.1093/bioinformatics/btp187.

